# Building an atlas of mechanobiology: high-throughput contractility screen of 2418 kinase inhibitors in five primary human cell types reveals selective divergent responses among related cell types

**DOI:** 10.1101/2025.01.11.632556

**Authors:** Anton Shpak, Jeremy Wan, Ricky Huang, Enrico Cortes, Yao Wang, Robert Damoiseaux, Ivan Pushkarsky

## Abstract

Cellular mechanical forces play crucial roles in both normal physiology and disease, yet drug discovery efforts targeting mechanobiology have been limited in part by assumptions about the conservation of contractile pathways across cell types. Here, we present the first high-throughput contractility screen of an annotated kinase inhibitor library, evaluating 2,418 compounds across five primary human cell types using the FLECS (Fluorescent Elastomer Contractility Sensors) platform. Quantification of contractile responses revealed selective divergent responses among related cell types. Clustering analysis identified distinct mechanobiological profiles and novel pathway associations that challenge the assumption that contractile pathways are too highly conserved for selective targeting. This systematic approach supports wider adoption of mechanical phenotypic screening as a viable strategy for discovering cell-type specific contractile pathway modulators for a broad range of mechanically-driven disease indications.

## 1. Introduction

Working alongside the well-established importance of chemistry and biology in health, the physics of cellular behavior—known as mechanobiology—also plays a crucial role. All cells, as well as the tissues and organs they constitute, possess the intrinsic ability to generate mechanical force. This capability is housed within the molecular machinery of each individual cell and is governed by a wide variety of pathways, many of which remain poorly understood. At the single-cell level, mechanical forces are utilized in processes such as cell division^1^, anchoring, and motility^2^, while in more complex scenarios, these forces contribute to functions such as wound repair^3^. At the tissue level, mechanical forces facilitate essential physiological processes, such as the contractions seen in cardiac, smooth, and skeletal muscle systems, which impact organs such as the bladder, intestine, and others. Mechanobiology is, therefore, a fundamental and ubiquitous feature of biological systems throughout the body^4, 5^ **(Fig 1)**.

**Figure 1:**
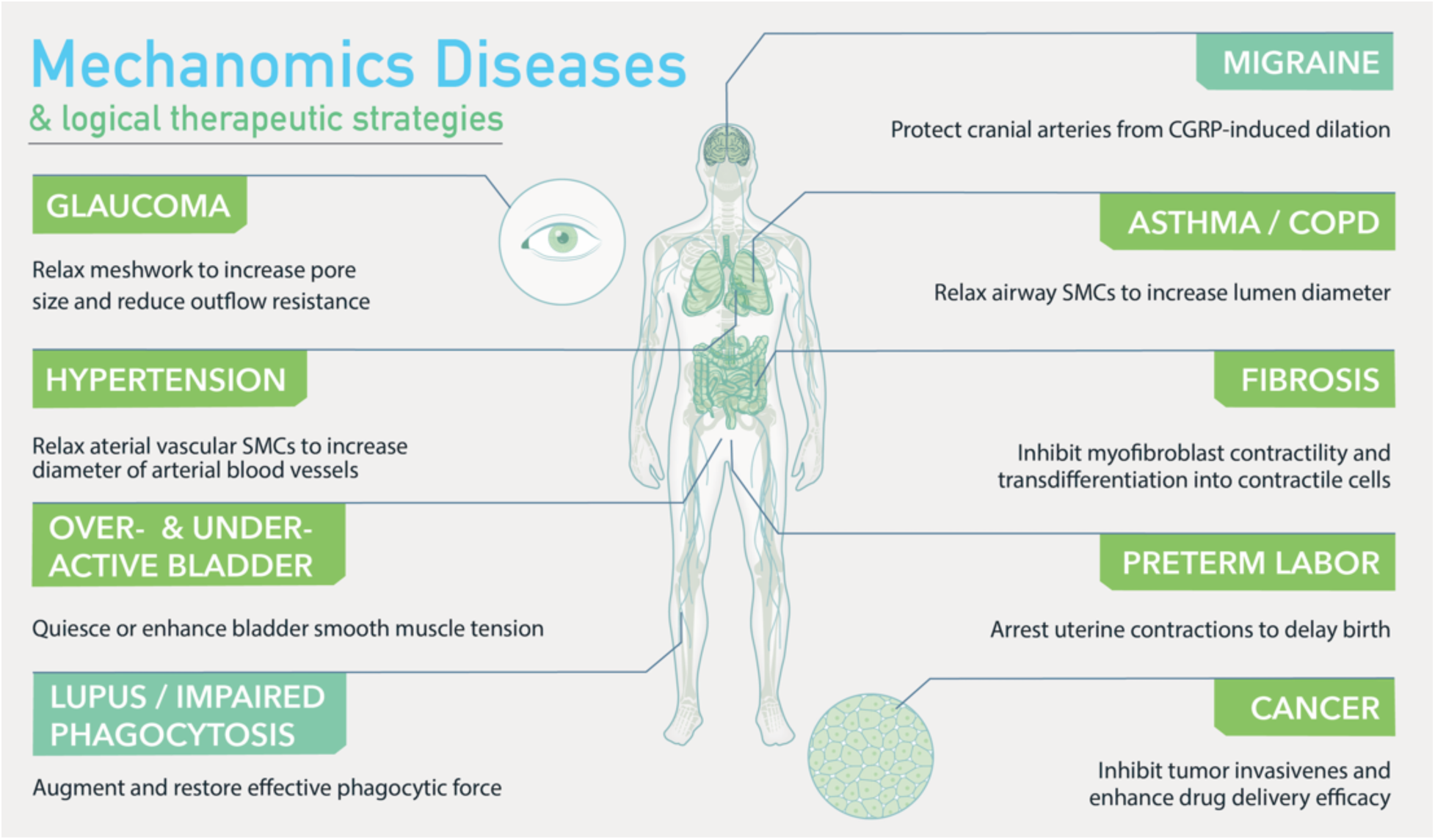
Body map of mechanobiological disease caused in part or in whole by contractile dysfunction in cells. Three main classes of disease are represented here: connective tissue such as asthma and spastic or underactive bladder, fibrotic disease such as lung fibrosis, and oncology where tumor microenvironments, metastasis and tumor tension all play key pathological roles. There is a major need to develop therapeutics that modulate the mechanobiological function in cells to treat these classes of diseases.

Dysregulation of these mechanical processes within cells can contribute, either partially or entirely, to the development of various complex diseases. These conditions can be grouped into several broad categories, including diseases related to connective tissue such as asthma^6–8^ and bladder dysfunction^9–11^, fibrotic conditions such as lung and liver fibrosis^12–14^, as well as oncological and rarer disorders. The development of therapies to target these diseases has been constrained by our limited understanding of the underlying biological mechanisms. There exist some therapeutic strategies that directly modulate contractile forces to treat disease. For instance, beta-2 adrenergic receptor agonists such as albuterol treat asthma by relaxing smooth muscle. However, overall, we lack comprehensive insight into the full spectrum of druggable pathways that regulate cellular and tissue mechanics.

Historically, a lack of technological means to quantitatively profile contractile cell force at sufficient scale, and the long-held conservative assumption that contractile pathways are highly conserved across cell types (making them challenging to target with precision and safety), has prevented a deeper pursuit of mechanobiological readouts as a means for phenotypic drug discovery.

To address this, we conducted the first ever high-throughput cell contractility screen of an annotated library of kinase inhibitors across two distinct disease categories within mechanobiology and utilizing five primary human cell types representative of these conditions. Our results reveal that, while some responses are conserved, there are also highly specific and selective activities observed across different cell types in response to the same treatment. These findings provide empirical evidence that selective pathways can indeed be exploited to safely target mechanical processes in the treatment of disease.

## 2. Results

### 2.1 FLECS technology is a general purpose platform to scalably screen contractile cell force

Previously, we introduced the FLECS Technology platform as an automated, high-throughput system for assaying contractile force at the single-cell level across large populations of various types of primary human cells^6, 8, 13, 15–19^. Briefly, this platform enables precise measurement of both acute and longer-term contractile responses in distinct cell types. Specifically, we have demonstrated its ability to classify the acute contractile responses of smooth muscle cells as well as the longer-term response of fibroblasts to TGF-β activation.

To explore how different human cell types and classes respond to the same pharmacological treatments, we screened a library of kinase inhibitors in two types of smooth muscle cells relevant to connective tissue disorders and two progenitor myofibroblast cell types involved in organ fibrosis. The smooth muscle cells, derived from bladder and airway tissues, were screened for their acute responses (45 minutes and 2 hours post-drug exposure) since conditions like asthma and spastic bladder require rapid intervention during flare-ups. The fibroblast progenitor cells, specifically lung fibroblasts and hepatic stellate cells, were screened for their responses over a 24-hour period in the presence of TGF-β, as this more closely models the activation of myofibroblasts—a key process in the progression of fibrosis.

### 2.2 Primary screen results

Following protocols developed in our prior works, we conducted acute response screens in primary human bladder (HBSM) and airway (HASM) smooth muscle cells (collectively, “SMC”), and 24-hour screens in primary human lung fibroblasts (HLF), hepatic stellate cells (HHSteC), and IPF patient derived lung fibroblasts (IPF-HLF) – collectively referred to as myofibroblast progenitors (“MYO”), each while exposed to TGF-β over this 24-hour period. Our experimental parameters are summarized in **Table 1**.

**Table 1.** Summary of the five primary human cell types and their screening conditions. SMC cells (blue rows) and MYO cells (orange rows) differ in assay format (acute vs. 24-hour), TGF-β activation, time points, and dosing. This table highlights how each cell type was tested under distinct protocols to capture disease-relevant contractile responses. The secondary 24hr screens in SMC were performed at the primary single-point level for cursory comparison to MYO responses, but these hits were not advanced into confirmation and subsequent stages.

| Cell Type | Abbrev. | Tissue / Origin | Screen Format | TGF-β? | Time Points | Dose(s) |
| --- | --- | --- | --- | --- | --- | --- |
| **Human Bladder Smooth Muscle**<br>("SMC") | HBSM | Primary bladder smooth muscle | **Acute** | No | Pre-drug, 45min & 2hr Post-drug | 35 nM, 356 nM |
|  |  |  | **Extended**<br>(secondary) | No | 24hr | 356 nM |
| **Human Airway Smooth Muscle**<br>("SMC") | HASM | Primary airway smooth muscle | **Acute** | No | Pre-drug, 45 min & 2hr post-drug | 35 nM, 356 nM |
|  |  |  | **Extended**<br>(secondary) | No | 24hr | 356 nM |
| **Lung Fibroblasts**<br>("MYO") | HLF | Primary human lung fibroblasts | **24-hour**<br>(activation) | Yes | 24hr | 35 nM, 356 nM |
| **IPF Lung Fibroblasts**<br>("MYO") | IPF-HLF | Patient-derived lung fibroblasts | **24-hour**<br>(activation) | Yes | 24hr | 35 nM, 356 nM |
| **Hepatic Stellate Cells**<br>("MYO") | HHSteC | Liver-resident stellate cells | **24-hour**<br>(activation) | Yes | 24hr | 35 nM, 356 nM |

All experiments were performed on 384-well FLECS plates at two doses (356 nM and 35 nM). In addition, to evaluate possible differential responses between the two categories of cell types, smooth muscle cells were also screened in a 24-hour format but without exposure to TGF-β at a single dose (356 nM). Collectively, over 38,000 drug-dosed wells were imaged each with >100 single cells, not including controls.

Acute SMC screens were performed by allowing seeded SMCs to adhere overnight on FLECS assay plates, imaging their contracted states at baseline, administering drug compounds, and reading their response at 45 minutes following drug addition. Final timepoints were taken 2 hours after drug administration. In this format, DMSO-only wells served as negative controls. No pharmacological positive control was administered since micropatterns lacking adhered cells serve as inherent positive controls.

All 24-hour SMC screens were conducted identically but were only imaged at 24 hours after drug addition. All 24-hour TGF-β MYO screens were performed by allowing MYO cells to adhere in FLECS assay plates pre-loaded with drug compounds for 4 hours. Then, TGF-β was added to the cells. After 24 hours, the plates were imaged at one terminal timepoint. In this format, DMSO-only wells served as negative controls while wells without TGF-β provided positive controls representing non-activated myofibroblast progenitor cells.

**Figure 3** shows the normalized screen results across all primary screens. Normalized contraction values for 24-hour screens are shown in Supplemental Information **(Fig S1)**. The tightest distributions are observed in acute SMC screens (hit upper bound >8 MADs below median of treated wells) while the most variation is seen in HLF at the 356 nM dose (hit upper bound 1.5 MADs below median). Robust Z’ factors for all TGF-β MYO screens are displayed in Supplemental Information **(Fig S2)**. For MYO screens, hits were defined as compounds that achieved both Z-score <-3 and normalized activity inhibition of at least 15% to ensure both statistical significance and biological relevance. For SMC screens, hits were defined as compounds that achieved a decrease in normalized activity inhibition of at least 20% from the baseline timepoint.

**Figure 2:**
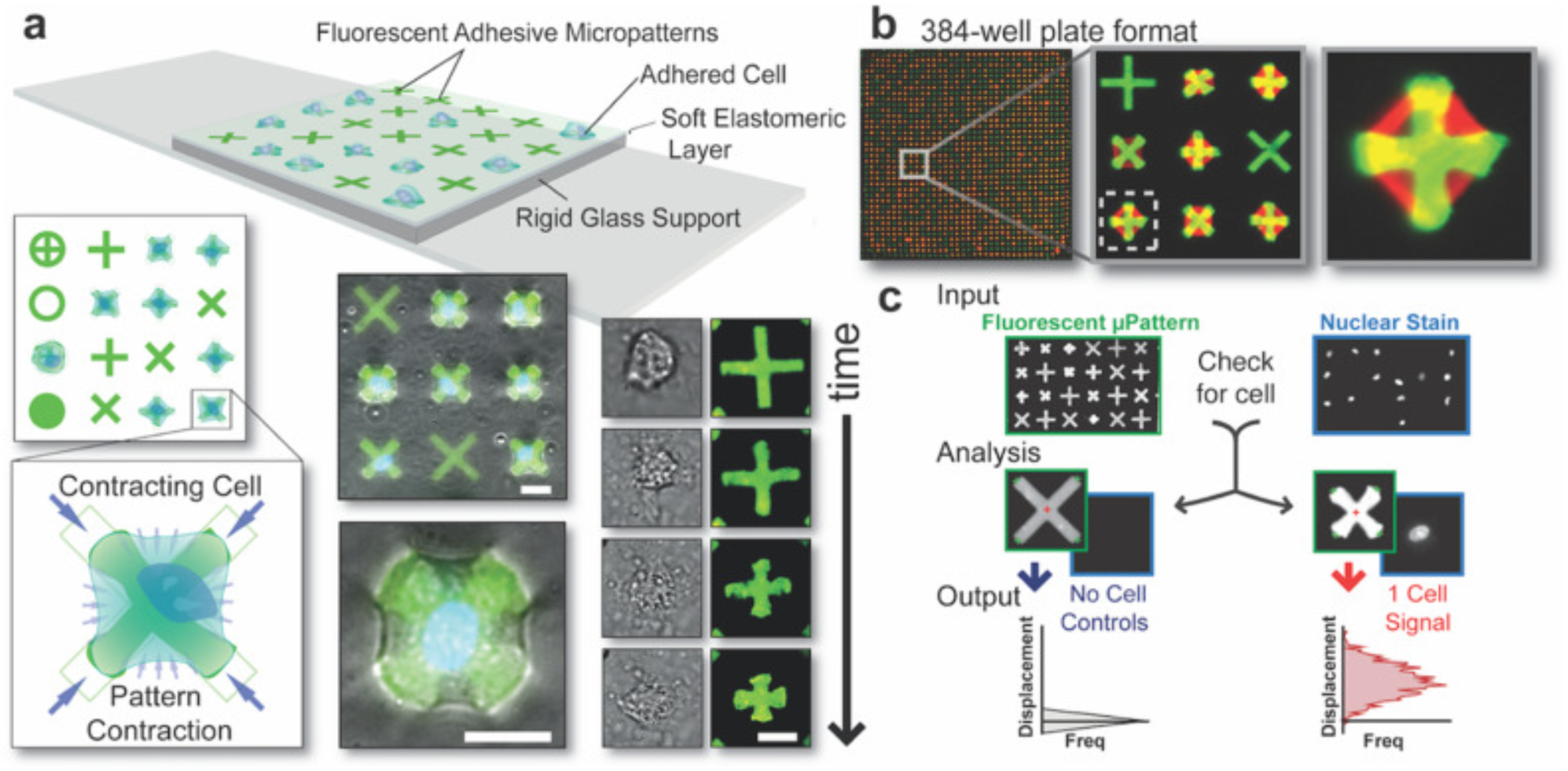
***(a)*** Schematic representation of the FLECS technology: Single cells adhered to adhesive micropatterns embedded within an elastomeric film exert mechanical force onto the micropatterns resulting in their displacements. Top view demonstrates various pattern shapes, while a magnified image illustrates a contracting cell inwardly displacing its terminals. Overlay of fluorescent patterns and phase contrast images captures contracting cells, with time-lapsed images showcasing the cell’s contraction over the underlying micropattern. Scale bars represent 25 μm. ***(b)*** Implementation of FLECS Technology in a 384 well-plate format. ***(c)*** Workflow for image analysis: Input includes aligned image sets of the micropatterns (set 1) and stained cell nuclei (set 2). Analysis involves (i) identification and measurement of all micropatterns in image set 1, (ii) cross-referencing the positions of each micropattern in image set 2, and (iii) determining the presence of 0, 1, or >2 nuclei (cells). Output consists of mean center-to-terminal displacements of micropatterns containing a single nucleus (one cell), compared to the median measurement of non-displaced patterns with zero nuclei. The resulting differences are presented as a horizontal histogram. MATLAB code enabling describing original versions of these algorithms are described in a previous work[16]. Figure adapted with permission from Pushkarsky et al. [13,15,16]

**Figure 3:**
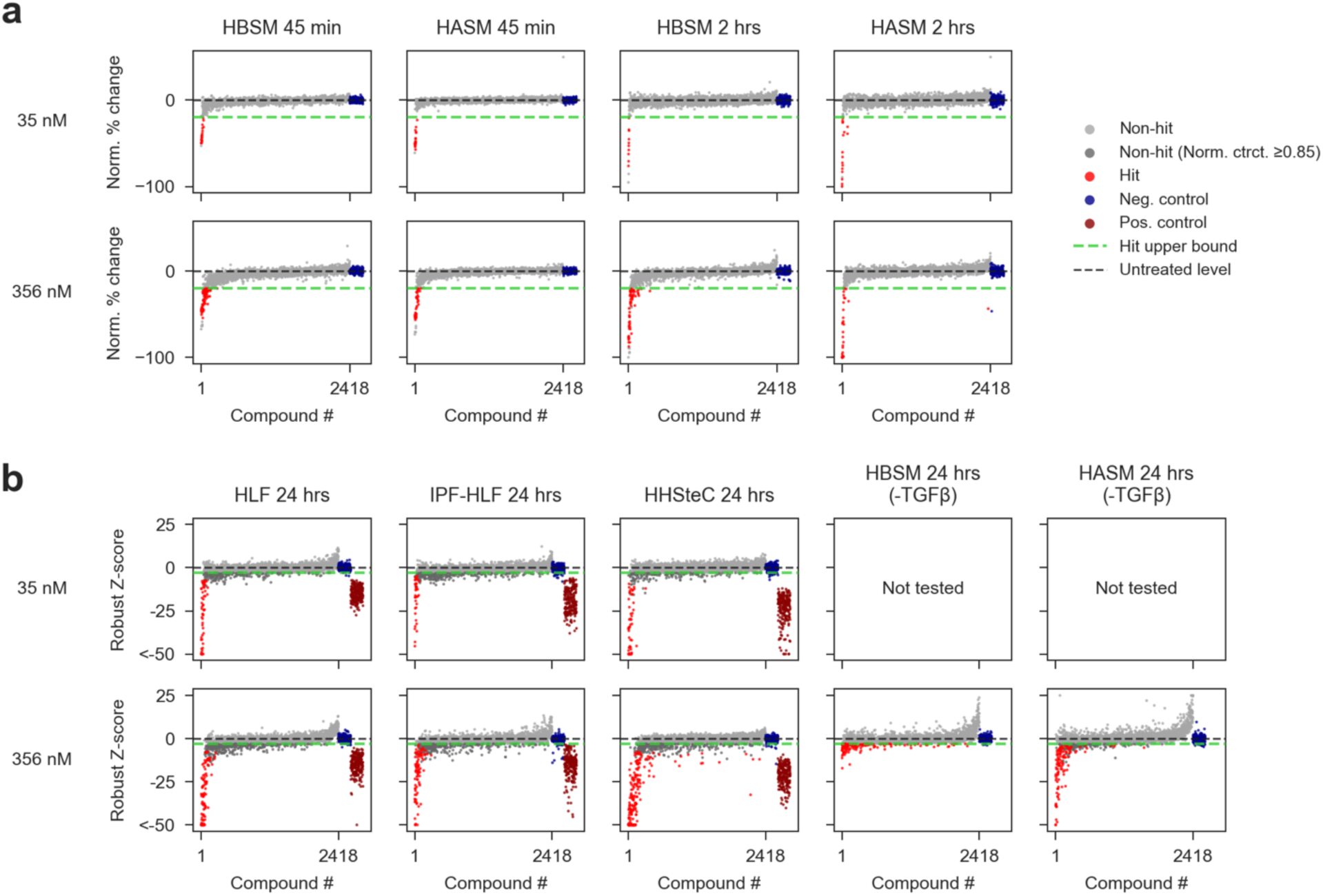
Results from primary contraction screens across multiple cell types. The data demonstrates the platform’s scalability and robustness, with clear separation between controls and hits. Expectedly, most compounds did not elicit significant responses, yet a notable subset of inhibitors was identified across cell types and doses. This underscores the non-random distribution of responses. A “Hit upper bound” threshold is marked by the green line, with non-hits below this threshold shaded in dark grey. These non-hits fail to meet normalized contraction constraints (<0.85) despite achieving robust Z-score criteria. ***(a)*** Scatter plots of acute SMC screen results. Robust Z-scores of contraction are shown, stratified by dose and cell type. Compounds are ordered by the average Z-score of contraction across all cell types and doses, with the same order maintained across all subplots. ***(b)*** Scatter plots of MYO screen results. Normalized percent change in contraction is shown, stratified by dose, timepoint, and cell type. Compounds are ordered by the average normalized percent change in contraction across all cell types, timepoints, and doses, with the same order maintained across all subplots.

### 2.3 Clustering of primary screening data identifies clear activity clusters described by specific pathway representation

To analyze broad patterns in compound responses, we applied agglomerative hierarchical clustering to the vectorized dataset **(Fig 4a)**, capturing compound activity across cell types and doses at the 24 hour timepoint.

**Figure 4:**
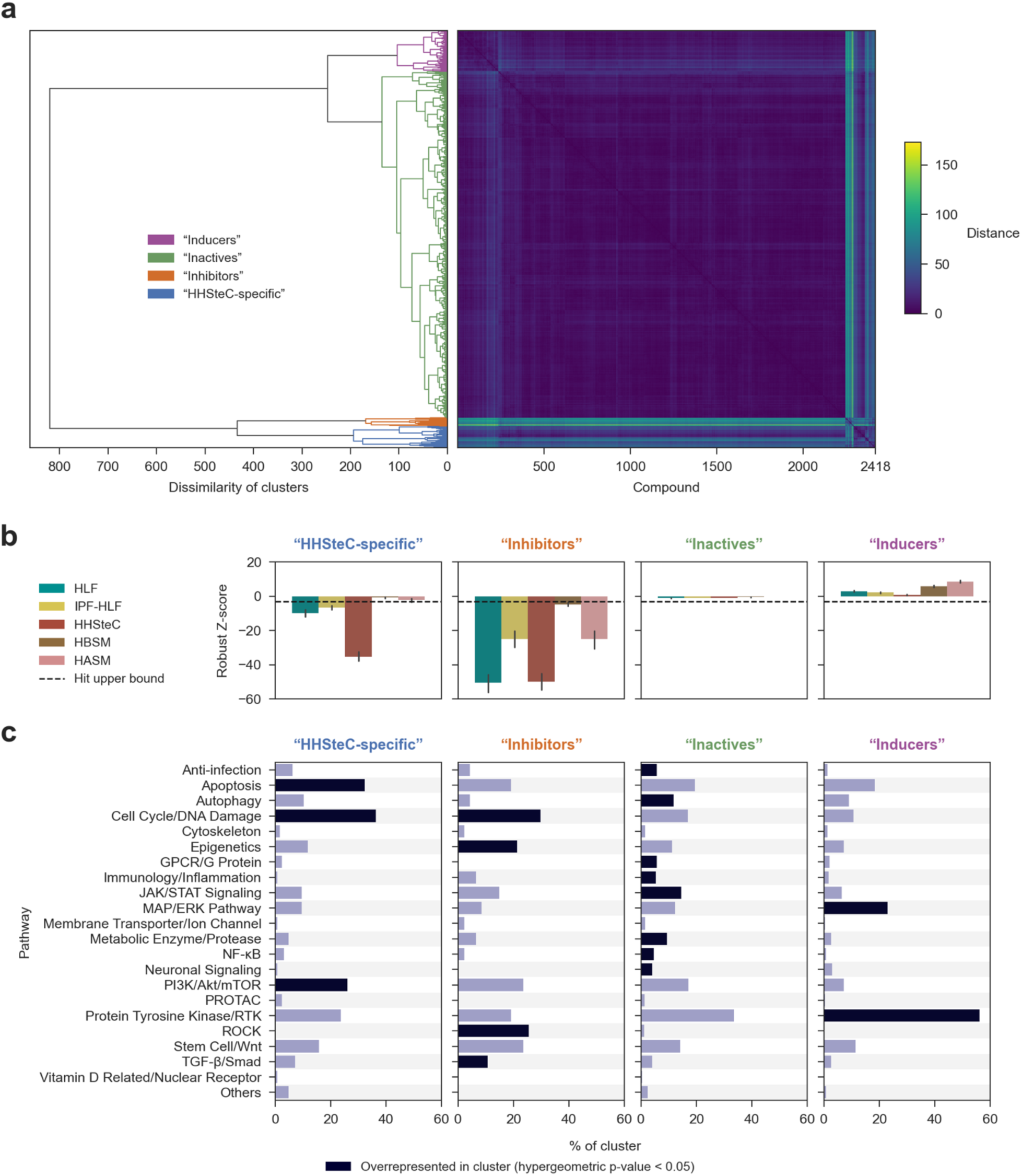
Clustering of 24-hour primary screen compounds reveals distinct contractile response types and pathway associations. ***(a)*** Dendrogram and pairwise distance matrix showing hierarchical clustering of compounds based on Z-score vectors across five cell types. Clusters highlight compounds with distinct response profiles, with pairwise distances >50 indicating highly divergent responses. ***(b)*** Bar graph comparing mean robust Z-scores of contraction (±95% CI) for each cell type within clusters, with the dotted line marking the Z = -3 upper bound for hits. ***(c)*** Distribution of compounds by affected pathways within each cluster, emphasizing pathway overrepresentation.

Agglomerative hierarchical clustering begins by treating each observation as its own cluster. Then, similar pairs of clusters are iteratively merged based on a linkage criterion that represents the dissimilarity between sets of observations. This criterion is computed as a function of the pairwise distances between the single observations within these sets.

Given the low dimensionality of our data, and given that magnitude of response is of primary concern, we used Euclidean distance as the pairwise distance metric. We performed the clustering using Ward’s linkage, which minimizes the increase in total within-cluster variance when merging two clusters. To visualize this process, we created a hierarchical dendrogram that represents relationships at various levels of similarity.

For this analysis, we focused on the penultimate branch point, where the dendrogram resolves into four clusters **(Fig 4a)**. We chose this number of clusters based on visual inspection. At the four-cluster level, the clusters appear sparse and well-separated, suggesting that they represent distinct and meaningful groupings. Moreover, the clusters remain large enough to yield statistically significant comparisons between them.

By examining the four primary clusters at this level, we identified fundamental groupings of compounds with similar mechanobiological effects, capturing both shared mechanisms and distinct response profiles across the dataset. Based on this analysis, we designated these clusters with the following labels:

1. *HHSteC-specific compounds* – compounds primarily active in human hepatic stellate cells (HHSteCs)
2. *Inhibitors* – compounds with broad inhibition across all cell types tested
3. *Inactives* – compounds that did not affect contraction in any cell type
4. *Inducers* – compounds that increased contraction of cells

Following from this, we used the hypergeometric distribution to evaluate which pathways were overrepresented within each cluster (under the null hypothesis that pathways were identically distributed across clusters). We identified several expected associations as well as novel ones. Specifically, the *Inhibitors* cluster contained overrepresentations of the cell-cycle/DNA damage, ROCK, and TGF-β/Smad pathways, all of which are known to affect cytoskeletal function, contraction, or TGF-β activation (which was induced in all MYO cell types). This result validated the FLECS platform’s ability to reflect known mechanobiological responses in cells.

Novel results from this analysis include the overrepresentation of PI3K/Akt/mTOR in the *HHSteC-specific* cluster but not in the *Inhibitors* cluster, and the overrepresentation of MAP/ERK in the *Inducers* cluster, suggesting possible targets for disease indications where enhancement or restoration of higher contractile tone would benefit patients, such as underactive bladder.

### 2.4 Subcluster dissection reveals distinct mechanobiological response profiles and pathway biases

To further refine our understanding of pathway groupings within the identified primary clusters, we performed a secondary clustering analysis. Each of the four main clusters—*HHSteC-specific compounds, Inhibitors, Inactives, and Inducers*—was further subdivided. The number of sub-clusters was chosen based on visual inspection of the dendrogram. This was done separately for each cluster, to ensure that sub-clusters were well-separated and were not singletons or otherwise too small to observe statistically significant differences. Overall, the subclusters maintained the general activity themes of the parent clusters. However, at this finer level of resolution, by similarly evaluating pathway overrepresentation using the hypergeometric distribution, additional mechanistic themes and pathway contributions that were not apparent at the primary cluster level were revealed.

In particular, the *HHSteC-specific* cluster was sub-divided into strong HHSteC inhibitors (HHS2), moderated HHSteC inhibitors (HHS3), and TGF-β pathway inhibitors which only affected MYO cells (HHS1 - where HLF, IPF-HLF, and HHSteC but not SMC cells were affected). Statistically significant overrepresentation of TGF-β pathway in HHS1 confirms this association.

The *Inhibitors* cluster broke out into strong general inhibitors (INH1), weak general inhibitors (INH2), and a third grouping which appears to represent strong inhibitors in non-diseased MYO cells but not in IPF-HLF nor the SMCs. Interestingly, INH1 (strong general inhibitors) are associated with metabolic enzyme pathway overrepresentation while INH2 (weak general inhibitors) are associated with ROCK pathway overrepresentation – a known broadly active regulator of cellular contractility.

The *Inactives* cluster—the largest by compound count—further resolved into three distinct groups: a wholly inactive group that accounted for over 75% of the parent cluster (INA1), as expected, and two smaller subclusters with unexpected low, non-zero activity one of which showed selective effects in HHSteCs (INA2), and one which displayed low non-zero activity across all cell types (INA3). This result suggests that on a relative basis, the modest activities of sub-clusters INA2, INA3 were more numerically similar to entirely inactive compounds indicating their activities would be masked without this secondary subcluster analysis.

Finally, the *Inducers* parent cluster was subdivided into two subclusters representing inducer activity in all cell types (IND1) and selective inducer activity only in SMCs (IND2). Pathway overrepresentations within these sub-clusters suggest that the MAP/ERK pathway might regulate contractility in various different cell types while Protein Tyrosine Kinase/RTKs pathway might be selective towards SMCs.

### 2.5 Hit selection and confirmation

Next, we cherry-picked molecules that registered as hits per our original definitions (see **4.4**). Some compounds that met the hit definition were excluded due to evidence of toxicity (observed through dead staining and from morphology). Other compounds were excluded due to inactivity in one cell type when picked together (e.g., compounds active in HLF at the 35 nM dose but only active in IPF-HLF at the 365 nM dose were picked at 35 nM and excluded from the analysis for IPF-HLF). A few borderline hits were excluded due suspected inactivity in combination with plate limitations (e.g., for HHSteC, there were slightly too many hits to fit on one cherry-pick drug plate).

Compounds were cherry-picked for confirmation screening only their lowest active doses. **Figure 6** summarizes our effort in taking primary screen data, identifying and retesting suspected hits, and advancing certain interesting confirmed hits into dose-response testing. As part of this process, certain molecules that were initially found to be more active in one cell type over another were re-tested across a dose-range in multiple cell types to better understand their selective activity.

**Figure 5:**
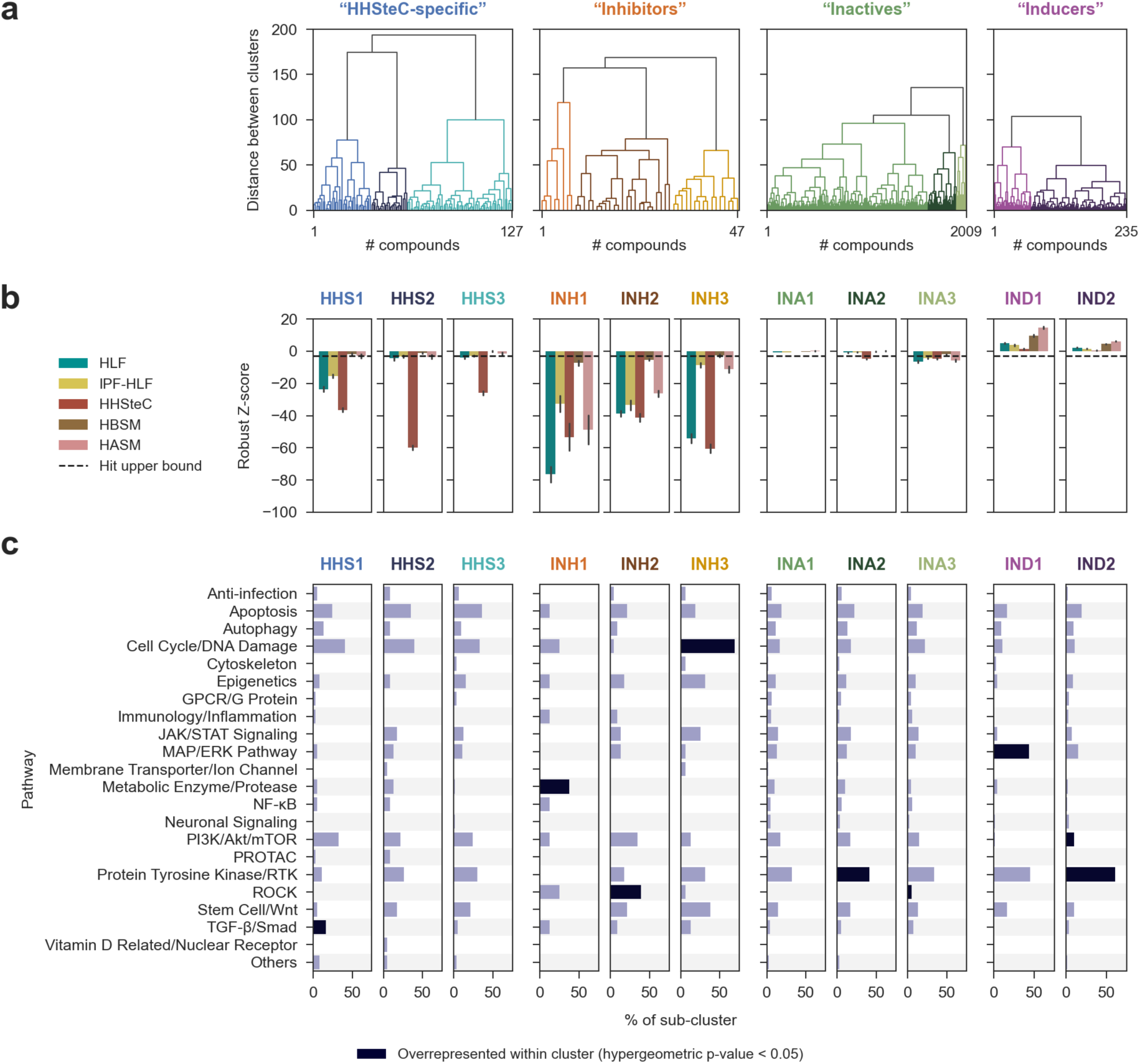
Sub-clustering of 24-hour primary screen compounds reveals distinct contractile response types and pathway associations. ***(a)*** Dendrogram and pairwise distance matrix showing hierarchical clustering of compounds based on Z-score. ***(b)*** Bar graph comparing mean robust Z-scores of contraction (±95% CI) for each cell type within sub-clusters, with the dotted line marking the Z = -3 upper bound for hits. ***(c)*** Distribution of compounds by affected pathways within each sub-cluster, emphasizing pathway overrepresentation within general cluster.

**Figure 6:**
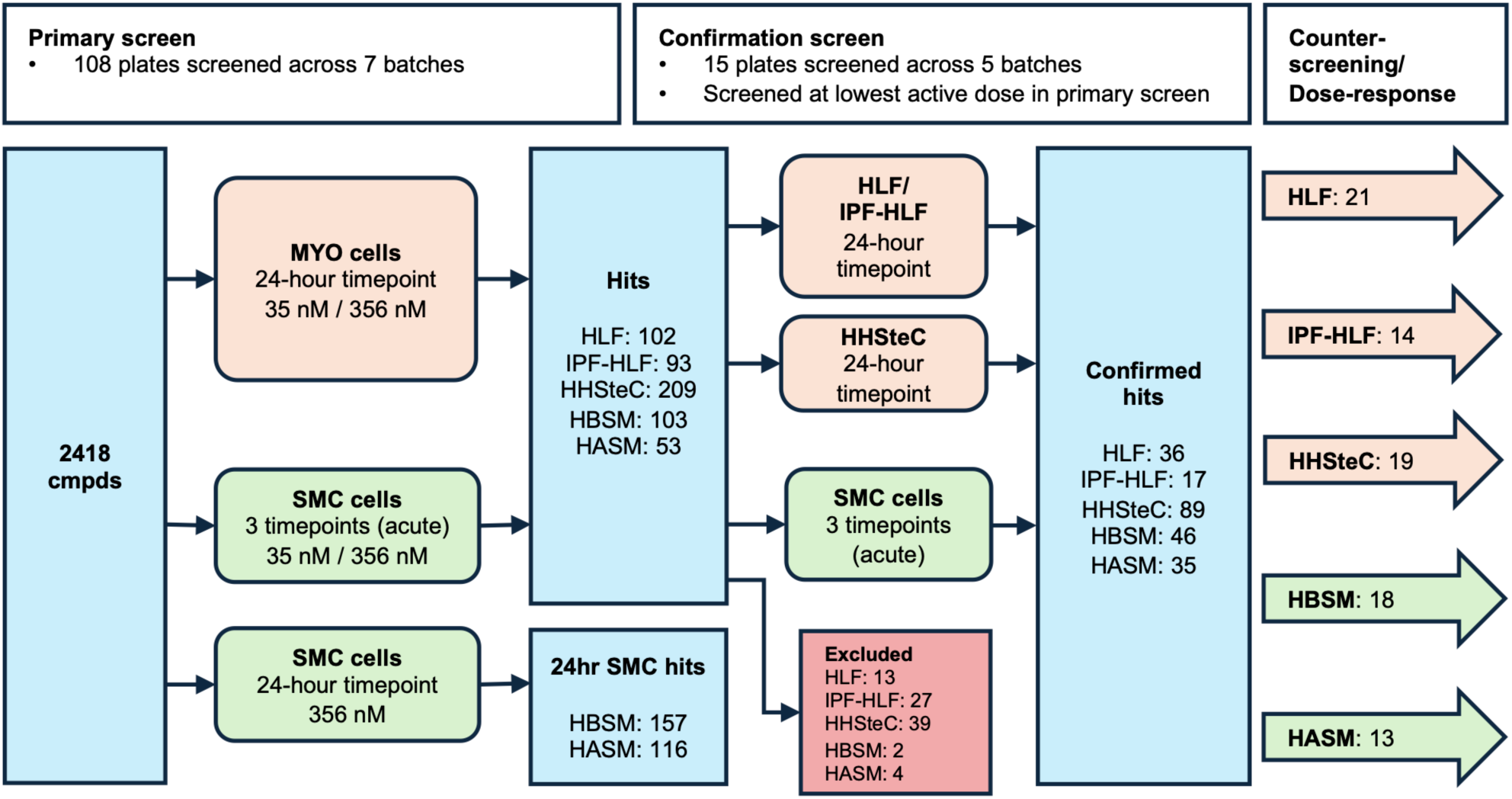
Screening flow-chart from primary screening to confirmation and dose response. Flowchart illustrating the screening process and the number of hits identified at each stage. A total of 2,418 compounds were screened at multiple doses in both MYO and SMC cell types, with imaging performed at biologically relevant time points. MYO cells were imaged at a single 24-hour time point to capture their response to TGF-β, which aligns with the biological timescale of myofibroblast activation. In contrast, SMCs were imaged pre-drug exposure, and at 45 minutes and 2 hours post-drug exposure to capture their acute responses. Hit definitions were established based on these time points. For comparison, SMCs were also screened at 24 hours; however, hits from this time point were not selected for further analysis as the focus remained on acute responses. Confirmation screens were subsequently conducted in triplicate using the same formats as the primary screen. Confirmed hits were defined as compounds which maintained their initial activity levels in all three of the triplicate tests.

As discussed previously, the SMCs, derived from bladder and airway tissues, were screened for their acute responses at 45 minutes and 2 hours post-drug exposure, as conditions like asthma and spastic bladder require rapid intervention during flare-ups. In contrast, the MYO cells— specifically lung fibroblasts and hepatic stellate cells—were screened for their responses over a 24-hour period in the presence of TGF-β. This longer time frame was chosen to model the activation of myofibroblasts, a key process in the progression of fibrosis. All subsequent confirmation screen analyses were conducted using these time-point-specific activity definitions.

Based on the strictest criteria of having all three replicates of the confirmation screen meet our definition for a hit, we observed a confirmation rate of >40% in all cell types with the exception of IPF-HLF which are primary IPF patient-derived and appear much more heterogenous in morphology, size, and contractile activity.

Data from the triplicate confirmation screen experiments demonstrated strong consistency, with mean pairwise R² correlation values exceeding 0.75 and approaching 0.9 for certain cell types **(Fig 7b)**, underscoring the robustness of the results. A heatmap of the number of hits (out of three replicates) across cell type and compound combinations **(Fig 7c)** reinforced our primary screening findings, revealing both highly cell-type-specific responses and shared effects across multiple cell types induced by certain compounds.

**Figure 7:**
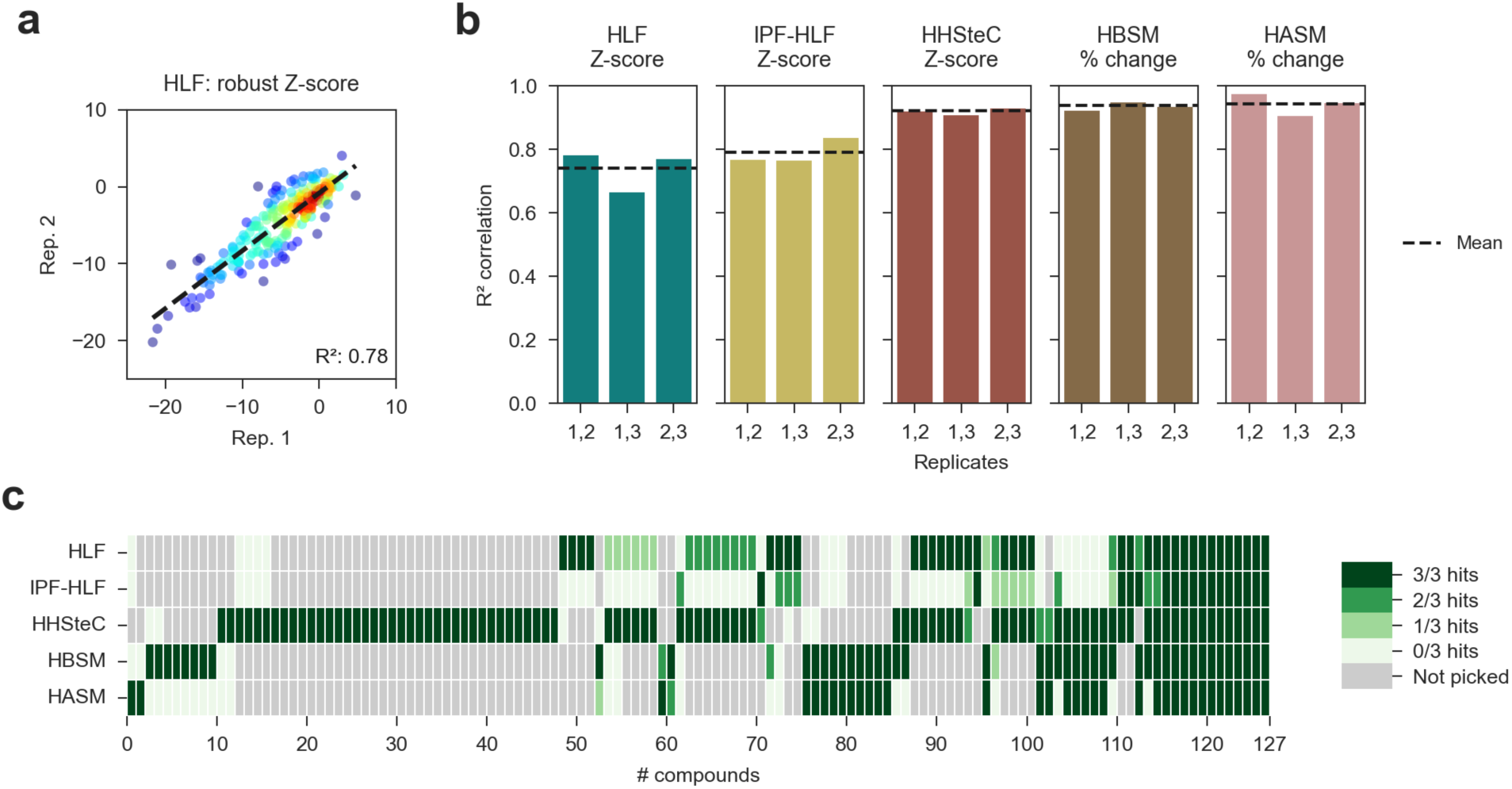
Confirmation screen results show strong correlation and cell-specific responses. ***(a)*** Scatter plot showing comparing robust Z-score of contraction between two replicates in HLF confirmation screen, colored by density. ***(b)*** Bar graphs showing correlation in response variables by cell type. Note that the SMC graphs show only the 45-min results. ***(c)*** Heatmap showing confirmation index by cell type for each compound. Note that some compounds were picked at different doses in different cell types. For SMC cells, the maximum number of hits across both timepoints is shown.

To further classify these diverse activities and elucidate the underlying biological pathways driving them, we performed agglomerative hierarchical clustering on the confirmation dataset and evaluated the pathways associated with the compounds comprising each cluster.

### 2.6 Pathway analysis of confirmation data

As with the earlier analysis of the 24-hour primary screen dataset, the number of clusters was chosen based on visual inspection of the clustering dendrogram **(Fig 8a)**. At the four-cluster level, the branches of the dendrogram appear well-separated, suggesting that these clusters represent meaningful groupings.

**Figure 8:**
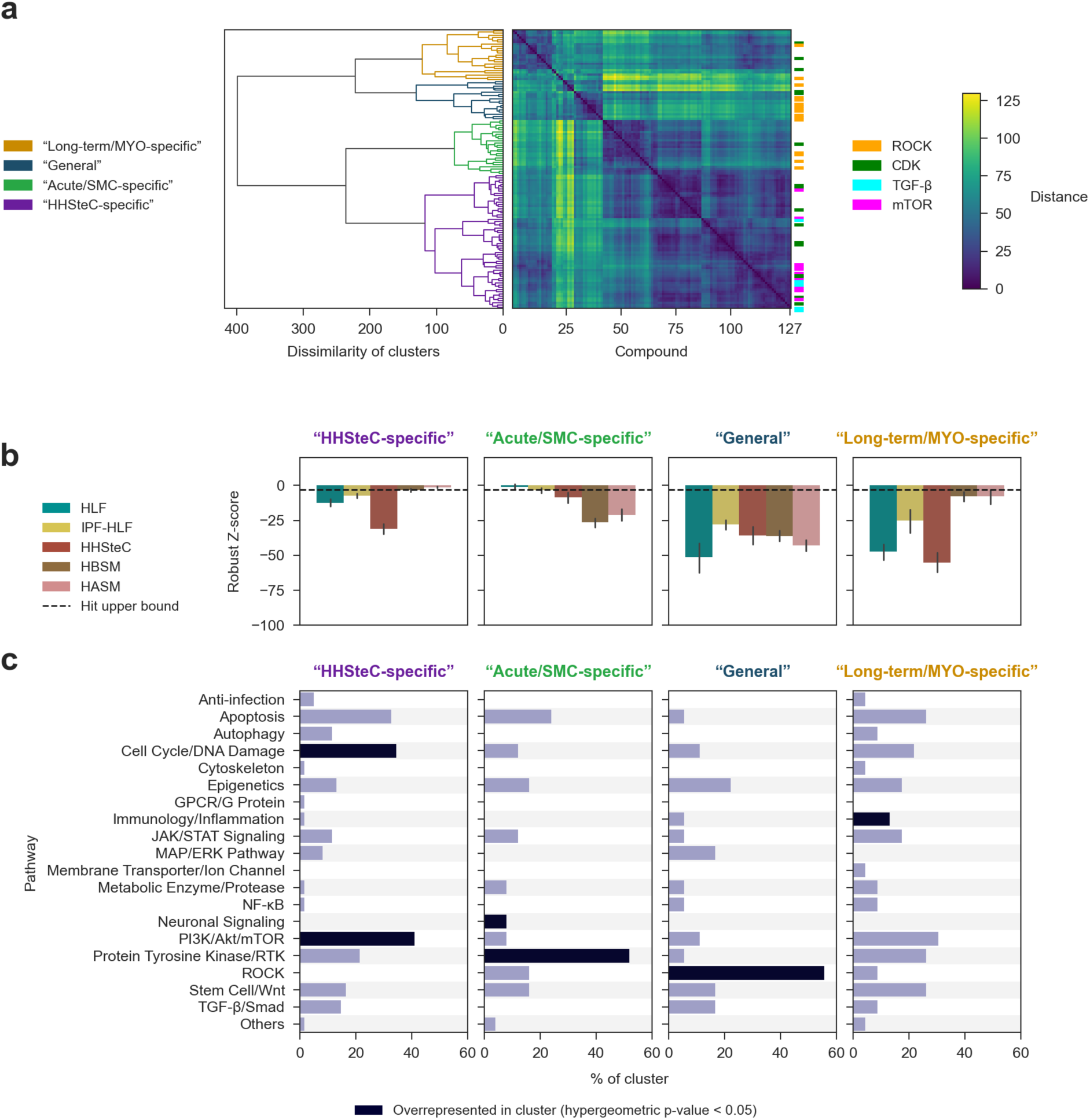
Clustering of confirmation screen results reveals distinct response types with differences in associated pathways. ***(a)*** Dendrogram and pairwise distance matrix showing hierarchical clustering of compounds based on Z-score vectors across five cell types. Selected compound targets are highlighted. ***(b)*** Bar graph comparing mean robust Z-scores of contraction (±95% CI) for each cell type within clusters, with the dotted line marking the Z = -3 upper bound for hits. ***(c)*** Distribution of compounds by affected pathways within each cluster, emphasizing pathway overrepresentation. Note: only 45-min data was included in activity vectors for SMC cells.

Within these four clusters, we identified fundamental groupings of compounds with similar mechanobiological effects, encompassing both shared mechanisms and distinct response profiles across the dataset. Two clusters characterized response types similar to those observed in the primary screen data – the *HHSteC-specific* cluster (typified by a greater response in HHSteC cells), and the *General* cluster (typified by a broad inhibitory response across all cell types), resemble the *HHSteC-specific* and *Inhibitors* primary screen clusters respectively. This suggests that these groupings reflect similarities in the constituent compounds across the two screens. Furthermore, cell-type-specific activities were observed in the remaining two clusters, which we designated as *Long-term/MYO-specific* and *Acute/SMC-specific*, reflecting the earlier observation time points used for SMCs.

To elucidate the pathways underpinning these activity profiles, we again applied the hypergeometric distribution to evaluate which pathways were overrepresented within each cluster (under the null hypothesis that pathways were identically distributed across clusters). Several expected associations were identified, along with novel ones **(Fig 8c)**. In particular, statistical overrepresentation of the ROCK pathway was observed in the *General* cluster, characterized by inhibition across all cell types. Meanwhile, the *Long-term/MYO-specific* cluster exhibited a statistical overrepresentation of immunology/inflammation pathways.

The PI3K/Akt/mTOR and Cell Cycle/DNA Damage pathways were again statistically overrepresented in the *HHSteC-specific* cluster, consistent with the analysis performed on the entire primary screen dataset. Interestingly, the Protein Tyrosine Kinase/RTK and Neuronal Signaling pathways were overrepresented in the *Acute/SMC-specific* cluster, suggesting a unique influence of these pathways on SMC contractile function or acute response times.

### 2.7 Comparison of HLF to HHSteC

Following our comparison of activities and profiles across different clusters, we next focused on analyzing potential differences in responses between related cell types, beginning with HLF and HHSteC, both classified as MYO cell types.

To do this, we first plotted the mean robust Z-scores (using confirmation screen data) for each compound tested in both cell types at the same dose against each other to directly compare their responses **(Fig 9a)**. Although the distribution of pathways represented among the confirmed hits was largely similar between the two cell types **(Fig 9b)**, the Z-score scatter plot suggests certain compounds were active in HHSteC but not HLF. This finding aligns with the presence of similar *HHSteC-specific* clusters observed in both the primary and confirmation screens.

**Figure 9:**
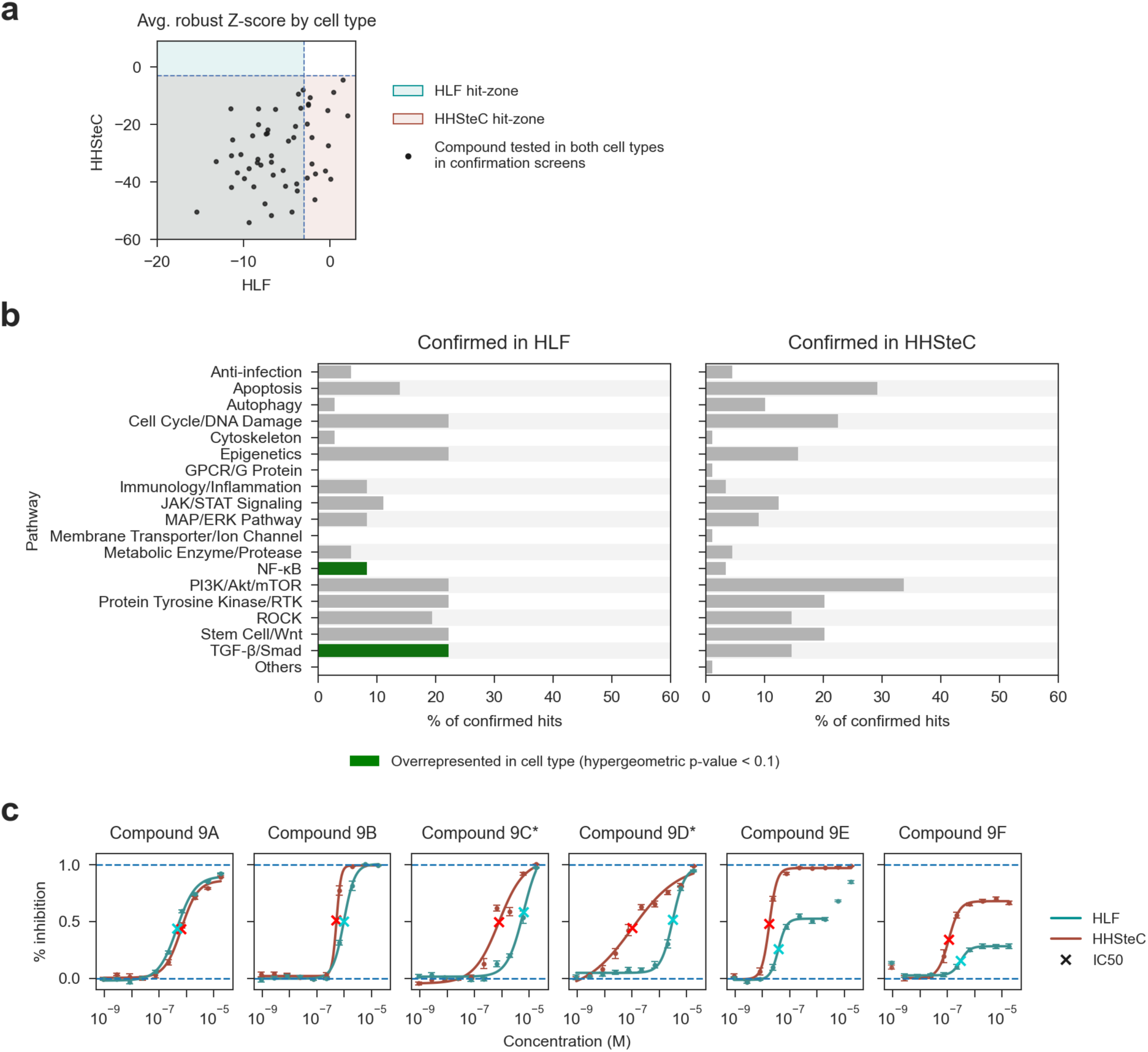
Comparing responses in HLF and HHSteC shows both general and selective responses. ***(a)*** Scatter plot of average Z-scores for contraction for compounds tested in both cell types at the same dose at the confirmation screen level. Note: not all hits were tested in both cell types, as in this case molecules were only re-tested in the cell type where they were originally identified as hits. ***(b)*** Distribution of compounds by affected pathways for confirmed hits in each cell type, highlighting pathway overrepresentation. ***(c)*** Dose response curves for selected compounds demonstrating cell-type selectivity among the MYO cells. Compounds labelled with an asterisk (*) were over 10 times more potent (by IC50) in HHSteC.

This analysis suggested the possibility of significant differential responses to the same treatments between these two related cell types. To test this hypothesis, we conducted concentration-response experiments on selected hits that appeared to exhibit such differences **(Fig 9c)**. Using the 24-hour MYO screening protocol, we evaluated a 10-step, 3-fold dose range spanning low nM to double-digit µM concentrations in both HLF and HHSteC.

Our findings revealed distinct responses in both potency and efficacy for certain compounds. For instance, compound 9A produced highly similar responses in the two cell types, with slightly better potency in HLF. In contrast, compounds 9C-9F exhibited marginally to significantly greater potency in HHSteC. Moreover, compounds 9E and 9F demonstrated over 100% greater percent inhibition in HHSteC compared to HLF at the response plateau level. Collectively, these dose-response curves demonstrate that highly related human cell types can exhibit substantially different responses to the same treatments under identical conditions.

### 2.8 Comparison of HASM to HBSM

Finally, we repeated this analysis on SMC cells (derived from bladder or airway tissue) used in the screens. As before, we plotted the mean robust Z-scores (from the confirmation screen) for each compound tested in both cell types at the same dose against each other to directly compare their responses **(Fig 10a)**.

**Figure 10:**
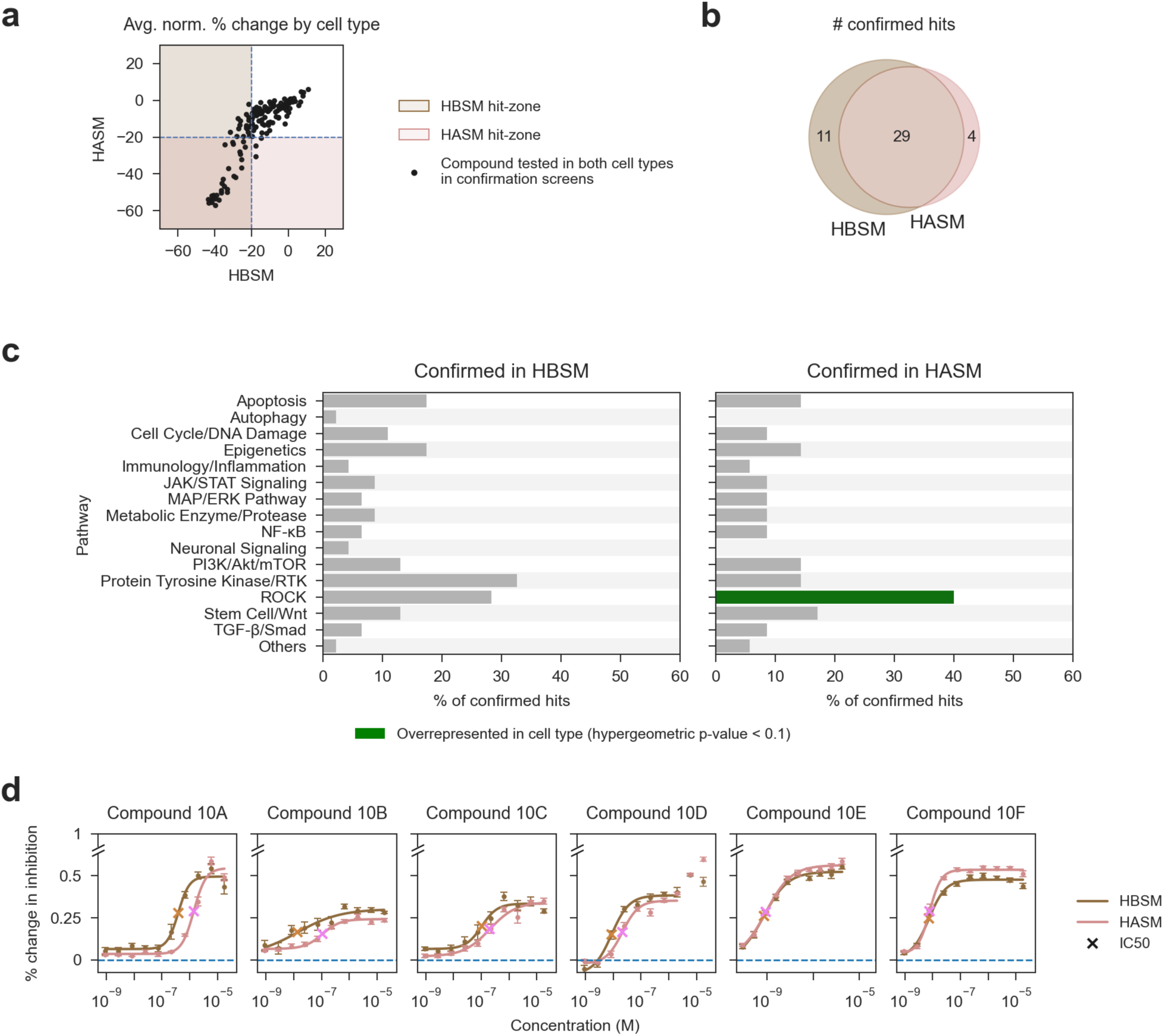
***(a)*** Scatter plot of average Z-score of contraction for compounds tested in both cell types at the same dose at the confirmation screen level. ***(b)*** Venn diagram showing overlap in confirmed hits. ***(c)*** Distribution of compounds by affected pathways for confirmed hits in each cell type, emphasizing pathway overrepresentation. ***(d)*** Dose response curves for selected compounds.

**Fig 10b** shows that the majority of confirmed hits were active in both cell types. However, some hits were not uniformly confirmed across all triplicates (e.g., not all three replicates met the criteria for a hit). This reflects the data presented in the Z-score scatter plot in which some hits are active but narrowly miss the hit criteria on an averaged basis. This suggests potential overall weaker activity in cases where only partial confirmation was achieved. Once again, the distribution of pathways represented among the confirmed hits were similar between the two cell types **(Fig 10c)**,

From this dataset, we selected six compounds for re-testing in both cell types under the same assay conditions as the original screen (i.e., 45-minute acute response to drug exposure) across a 10-step, 3-fold dose range spanning low nM to double-digit µM concentrations.

Although the differences were less pronounced than those observed in the MYO comparison **(Fig 9)**, we still identified distinct responses between the two cell types. Specifically, compounds 10A–10D exhibited better potency in HBSM with similar efficacy in both cell types. Conversely, compound 10F, and to a lesser extent 10E, demonstrated greater percent inhibition in HASM compared to HBSM.

Importantly, all experiments were conducted simultaneously on the same plates, ruling out batch effects or inter-experimental variability as potential causes for these differences. Collectively, these findings further demonstrate that highly related cell types, such as primary human bladder smooth muscle cells (HBSM) and primary human airway smooth muscle cells (HASM), can exhibit distinct responses to the same compound treatments.

## 3. Discussion

The purpose of this study was twofold.

First, to demonstrate that contractility and mechanical cell function, though governed by the same family of contractile proteins, can exhibit distinct regulatory mechanisms across cell types. Our findings, derived from the first-ever high-throughput kinase inhibitor screen targeting cell contraction, provide compelling evidence that mechanobiological pathways are diverse and selectively targetable. This suggests that direct screening for contraction modulators could unveil novel, first-in-class therapeutics with potential for high therapeutic indices, even among related cell or tissue types.

Second, this work highlights the unprecedented capability and scalability of the FLECS platform. For the first time, we were able to scalably quantify the effects of >2,400 annotated kinase inhibitors across five cell-types at multiple doses, as well as perform confirmation and dose-response studies. This was accomplished all using the same assay and readout, in under 8 weeks of run time, including plate manufacturing (in house) and data analysis. This dataset comprises over 38,000 unique drug-loaded wells each with 100s of single-cell datapoints in a single study. This scale and quantitative resolution enables a network level of insight into mechanobiology laying the groundwork for building a mechanobiology atlas, a resource with the potential to transform how we approach drug discovery and deepen our understanding of cellular mechanics.

Our systematic analysis of kinase inhibitors revealed that cellular contractility can be selectively modulated by small molecules, with even related cell types exhibiting divergent responses to the same compounds. While many compounds showed similar effects across cell types, we identified specific cases where responses clearly diverged. This divergence was particularly well-demonstrated in MYO cells, where the distinct response patterns were confirmed in dose-response experiments. SMCs also exhibited divergent responses, though less consistently when examined in targeted dose-response studies compared to the initial screens. Collectively, these results underscore the complexity of mechanobiological pathways and demonstrate that contractility regulation can differ even among related cell types, suggesting the possibility of selectively regulating contractile phenotypes with good therapeutic indices.

Our clustering analyses of the dataset revealed both expected and novel mechanobiological signatures. We identified known pathway associations like ROCK^20, 21^ and TGF-β, validating our approach, while also discovering unexpected pathway overrepresentations such as MAP/ERK in SMCs and PI3K/Akt/mTOR in HHSteCs, suggesting new therapeutic opportunities like enhancing contractility in underactive bladder through MAP/ERK modulation. Interestingly, despite using TGF-β for myofibroblast activation, the MYO-specific cluster did not show statistical overrepresentation of the TGF-β/SMAD pathway, likely because these compounds generated activity profiles that clustered with other mechanobiological response patterns rather than forming a distinct signature. This distribution of responses across multiple clusters suggests that mechanical responses to kinase inhibitors may not cleanly segregate by canonical pathway annotations. Moreover, these clustering patterns reveal the complex evolutionary landscape of contractile regulation, suggesting that while core contractile machinery is conserved, its regulation has diversified to meet tissue-specific functional demands.

The robustness of our findings is supported by high confirmation rates exceeding 40% across triplicates in most cell types, though we noted some important technical considerations. Lower confirmation rates in patient-derived IPF-HLF cells reflect the inherent variability of such samples, highlighting the importance of multiple cell models in translational research. Additionally, compound storage conditions may have affected the reproducibility of weakly active compounds, though this limitation did not impact our major hits, which showed consistent activity across experiments. Future studies will benefit from standardized compound preparation protocols across all screening stages.

The creation of this first mechanobiological response atlas represents a significant advance in our understanding of cellular mechanics. By systematically mapping how diverse cell types respond to pharmaceutical perturbations, we establish a framework for predicting tissue-specific drug effects on mechanical properties. This approach could revolutionize drug development by enabling early identification of both desired tissue-specific effects and potential mechanical side effects. The atlas concept could be further expanded by incorporating additional cell types, drug classes beyond kinase inhibitors, and complementary readouts such as transcriptional profiles. Such a comprehensive resource would provide unprecedented insight into the relationship between chemical structure, pathway activation, and mechanical outcomes across tissues.

In conclusion, this work advances our understanding of mechanobiology, demonstrating that contractility pathways are not universally conserved and that selective targeting is feasible. Our findings suggest new strategies for drug development, where mechanical phenotypes could be modulated with tissue specificity despite shared molecular machinery. It also showcases the transformative potential of the FLECS platform to uncover novel therapeutic targets and create a comprehensive mechanobiology atlas. These findings lay the groundwork for future studies and provide a strong foundation for translating these insights into meaningful therapeutic advances.

## 4. Methods

### 4.1 Cell culture

Cryopreserved human primary lung fibroblasts (HLF), human primary hepatic stellate cells (HHSteC), human airway smooth muscle (HASM) cells, and human bladder smooth muscle (HBSM) cells were obtained from ScienCell. IPF-HLF were a gift from Prof. Brigette Gomperts (UCLA). HLF and IPF-HLF were cultured in Lung Fibroblast Growth Medium (Cell Applications, catalog #516–500),). HHSteC were cultured in Stellate Cell Medium (SteCM, Cat. #5301). HASM and HBSM cells were maintained in Ham’s F-12 medium supplemented with 10% fetal bovine serum (FBS) and 1% penicillin/streptomycin. All cell types were grown in T75 flasks under standard conditions (37 °C, 5% CO₂) until ready for experiments. Cells were detached using 0.05% Trypsin-EDTA and prepared for experiments at passage 3. For experiments, HLF, HHSteC and IPF-HLF were tested in Dulbecco’s Modified Eagle Medium (DMEM) containing 10% FBS and 1% penicillin-streptomycin, while HASM and HBSM cells were tested in their respective Ham’s F-12 growth medium.

### 4.2 FLECS contractility assay protocol

The FLECS contractility assay followed protocols previously published^16^. Plates pre-coated with 70 µm “X”-shaped collagen type IV micropatterns on an 8 kPa substrate (Forcyte Biotechnologies, product #384-HC4R-QC10) were used throughout the study. Wells were initially filled with 25 µL of either serum-free DMEM (MYO cells) or Ham’s F12 media (SMC) supplemented with 1% penicillin-streptomycin. The appropriate cells were resuspended at 50,000 cells/mL in their proper culture medium. Cells were seeded in 25 µL volumes per well and allowed to settle for 1.5 hours at room temperature before being transferred to a 37 °C incubator. For MYO cells, compounds were added via Biomek automation 1.5 hours after incubation, followed by 10 minutes of plate shaking. TGF-β1 (PeproTech, 2.5 ng/mL) was added 30 minutes after compound addition. Plates were then returned to the incubator. At 24 hours post-TGF-β1 addition, Hoechst 33342 live nuclear stain (1 µg/mL) was added, and plates were imaged after 30 minutes using a Molecular Devices ImageXpress automated microscope. For SMC cells, plates were incubated for 24 hours post-seeding. Baseline images were acquired before compound addition via Biomek, followed by 10 minutes of plate shaking. Wells were then imaged at 45 minutes and 2 hours post-compound addition. Prior to the 2-hour imaging timepoint, Hoechst 33342 live nuclear stain was added for 30 minutes to enable nuclear imaging. Analysis of cell contraction and adhesion was performed using proprietary computer vision algorithms developed by Forcyte^16^.

### 4.3 Compound libraries

The screen was conducted using the Kinase Inhibitor Library (MedChemExpress), which consists of 18 unique 384-well source plates. Each source plate is laid out according to one of nine unique plate maps, and each map is repeated at two different stock concentrations.

### 4.4 Hit criteria

For MYO cells, compounds were selected as ‘hits’ for retesting if the robust Z-score of contraction was more than -3 (>3 MADs below the median of the negative controls) and the normalized inhibition was at least 15%. Primary screen hits were also filtered for toxicity based on morphology. For SMC cells, compounds were selected as ‘hits’ for retesting for a given timepoint if the normalized percent decrease in contraction (compared to the baseline timepoint) was greater than 20%. Primary screen hits were also filtered for toxicity based on dead stain imaging, with a cutoff of 3 times the median dead percent of the negative controls. Compounds were defined to be “confirmed hits” in a given cell type if after being chosen for retesting, they met the above conditions for 3/3 replicates. In addition, all preliminary hits were also filtered for toxicity based on dead stain imaging, with a cutoff of three times the median dead percent of the negative controls. If any of the three replicates were beyond this cutoff, compounds were excluded from the confirmed hits.

### 4.5 Data analysis

Hierarchical agglomerative clustering was performed using the AgglomerativeClustering() function from the scikit-learn Python library. Compounds were characterized by vectors with five components, with each component corresponding to the robust Z-score of contraction (for MYO cells) or the robust Z-score of the normalized percent inhibition (for SMC cells) in a given cell type. Clustering was done using Ward’s linkage, computed from the Euclidean distance metric. The number of clusters for comparison and the corresponding distance threshold in each case was chosen based on visual inspection of the clustering dendrogram. Clusters were chosen at the point where branches of the dendrogram appeared most sparse and well-separated, deviating where necessary to ensure that (a) there were no singletons/clusters too small to yield statistically significant differences and (b) meaningful differences in response were not obscured. Pathway overrepresentation was analyzed using the hypergeometric distribution. This was performed using the hypergeom.cdf() function from the SciPy Python library. The p-value that a pathway is overrepresented was computed as the probability that a given cluster (or group of confirmed hits for a given cell type) contains at least the observed number compounds affecting that pathway when drawing groups of the observed size randomly without replacement under the null hypothesis that pathways are evenly distributed throughout the population.

## Author Contribution

I.P. conceived of the project. E.C., Y.W., R.H., J.W., and A.S. performed the experiments. A.S., E.C., J.W., Y.W., and I.P. analyzed the data. R.D. developed the automation strategy and oversaw the robotic laboratory and lead compound management. A.S. developed data analysis and interpretation scripts and prepared the figures. A.S., I.P., Y.W., and R.D interpreted the data. I.P., A.S., and R.D. wrote the manuscript.

## Acknowledgements

The authors thank the Magnify Incubator at CNSI and the Nanolab at UCLA, and the Molecular Screening Shared Resource at CNSI for providing critical infrastructure to support the work. The authors also thank Mason Victors for his invaluable guidance regarding robust assay development and execution practices to ensure data quality and on relevant data analysis approaches.

