## Supplemental Figures S1-S3 for "Building an atlas of mechanobiology: high-throughput contractility screen of 2418 kinase inhibitors in five primary human cell types reveals selective divergent responses among related cell types"

**Authors & Affiliations**

Anton Shpak[a], Jeremy Wan[a], Ricky Huang[a], Enrico Cortes[a], Yao Wang[a], Robert Damoiseaux[a,b,c], Ivan Pushkarsky[a]*

*Corresponding author.

Authors 2–5 are listed in inverse order of the day of their birthday.

[a]Forcyte Biotechnologies, Inc

[b]University of California Los Angeles, Los Angeles, CA 90095, United States

[c]California NanoSystems Institute at UCLA, Los Angeles, CA 90095, United States

**Table of Contents**

Figure S1: Normalized contraction in 24-hour primary screens
Figure S2: Robust Z'-factor by cell type/screen batch
Figure S3: Comparing responses in HLF and IPF-HLF

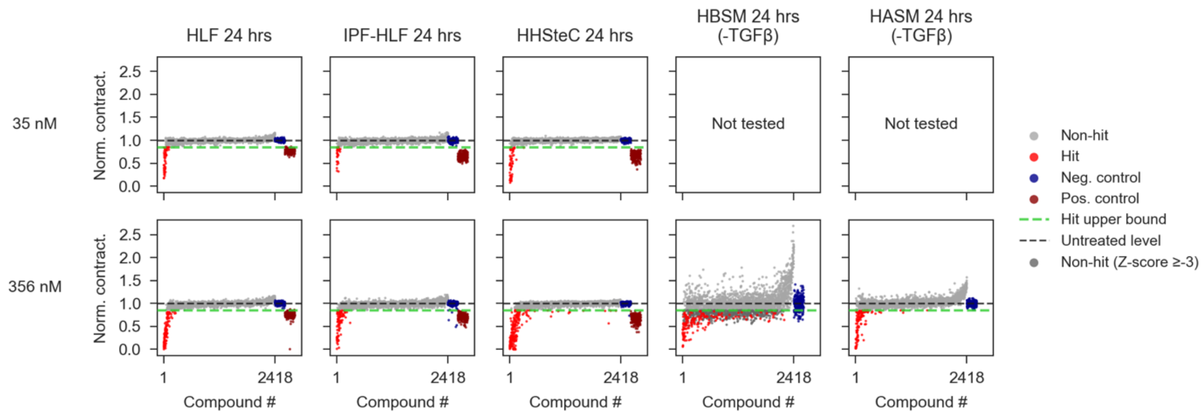


**Figure S1: Normalized contraction in 24-hour primary screens**

Scatter plots of 24-hour screen results. Normalized contraction values are shown, stratified by dose and cell type. Compounds are ordered by the average robust Z-score across all cell types, and doses (the same order as in **Fig 3b**), with the same order maintained across all subplots. A “Hit upper bound” threshold is marked by the green line, with non-hits below this threshold shaded in dark grey. These non-hits fail to meet robust Z-score constraints (<-3) despite achieving normalized contraction criteria. The results demonstrate that even when hits appear to be “borderline,” having Z-scores close to the Z = -3 cutoff, they generally fall well beyond the normalized inhibition constraint and thus represent a significant decrease in contraction.

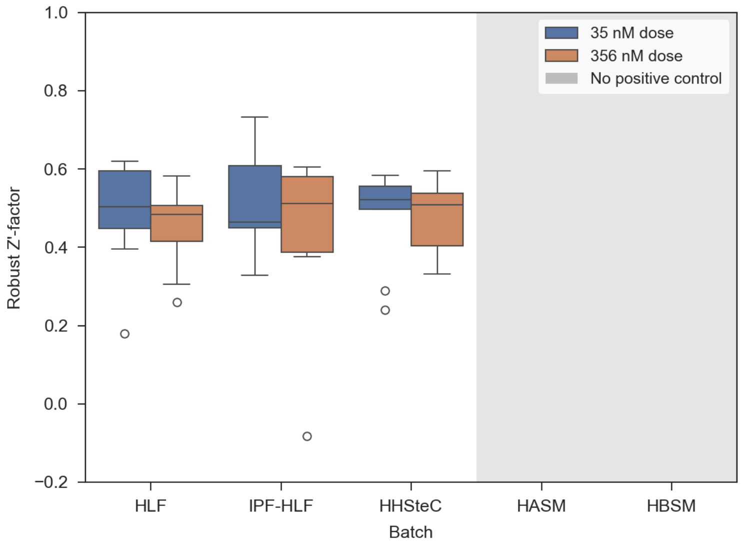


1

**Figure S2: Robust Z'-factor by cell type/screen batch**. Boxplot showing aggregated robust Z'-factors of each screen plate, separated by dose and cell type. The Z'-factors are generally around 0.5, indicating good separation of positive/negative controls in most plates screened. Z'-factors seem to be generally consistent across cell types and doses. This underscores the validity of the data and the robustness of the assay. Note that a few low outliers are observed. The robust Z'-factor was calculated as follows:

$$\text{Robust Z'} - \text{factor} = 1 - \frac{3\left(MAD_p + MAD_n\right)}{\left|\tilde{X}_p - \tilde{X}_n\right|}$$

where $MAD_p$, $MAD_n$, $\tilde{X}_p$, and $\tilde{X}_n$ are the median absolute deviation (MAD) of the positive controls, MAD of the negative controls, median of the positive controls, and median of the negative controls respectively. HASM and HBSM screens did not have a positive control and so this statistic is not applied in those cases.

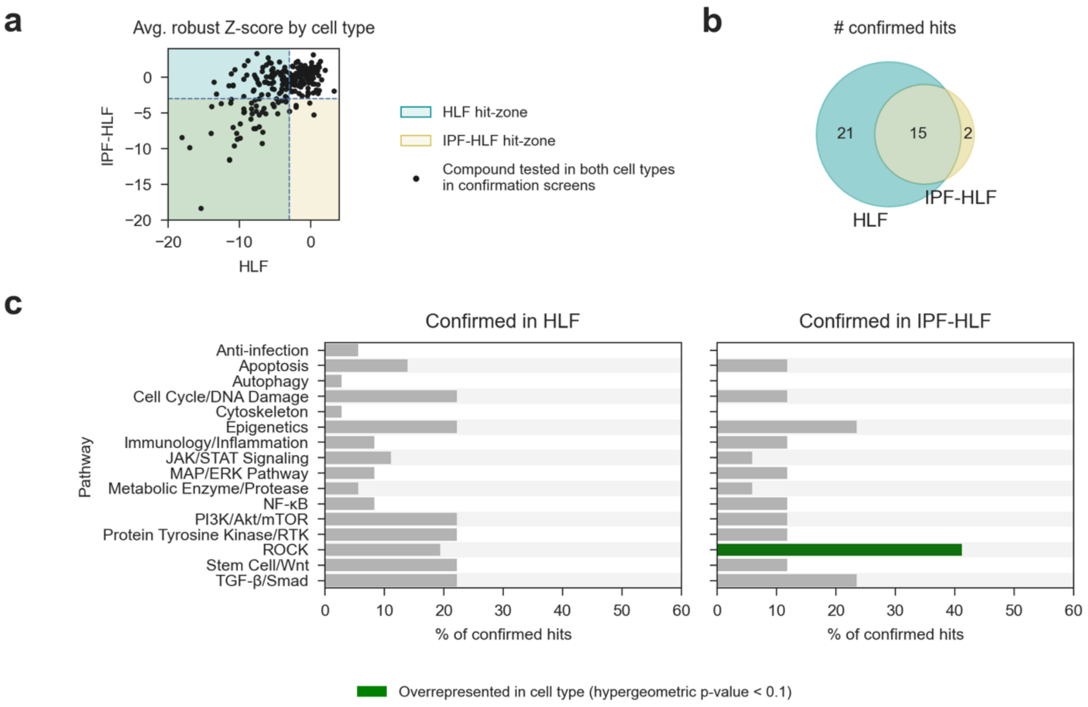


1

**Figure S3: Comparing responses in HLF and IPF-HLF shows both general and selective responses.** ***(a)*** Scatter plot of average Z-score of contraction for compounds tested in both cell types at the same dose at the confirmation screen level. ***(b)*** Venn diagram showing overlap in confirmed hits. ***(c)*** Distribution of compounds by affected pathways for confirmed hits in each cell type, emphasizing pathway overrepresentation. Although the general pathway distribution is similar, the ROCK pathway is overrepresented in IPF-HLF confirmed hits.
